# A PRISMA-guided systematic review of musculoskeletal modelling approaches in lower-limb cycling biomechanics

**DOI:** 10.64898/2026.03.05.709765

**Authors:** Ana C. C. de Sousa, Alexandre B. Peres, Josep M. Font-Llagunes, Roberto de S. Baptista, Rosa Pàmies-Vilà

**Affiliations:** School of Engineering - Instituto Químico de Sarriá (IQS), Universidad Ramon Llull (URL), Barcelona, Catalunya, Spain; Department of Mechanical Engineering and Institute for Research and Innovation in Health, Universitat Politècnica de Catalunya - BarcelonaTech (UPC), Barcelona, Catalunya, Spain; Department of Electrical Engineering - School of Technology, Universidade de Braśılia (UnB), Brasília, Distrito Federal, Brazil; Institut de Recerca Sant Joan de Déu, Esplugues de Llobregat, Catalunya, Spain

## Abstract

Cycling is commonly employed in sports performance, rehabilitation, and clinical contexts, while musculoskeletal (MSK) simulations enable the investigation of internal biomechanics that cannot be measured experimentally. Despite growing use, the application, validation, and standardisation of MSK simulations in cycling remain unclear. This review aimed to systematically characterise the application, validation strategies, modelling assumptions, and reporting practices of musculoskeletal simulations in lower-limb cycling biomechanics. Searches were performed in Scopus, PubMed, IEEE Xplore, and Web of Science on 1 August 2024, covering studies from January 2010 to July 2024. Peer-reviewed English-language journal articles applying MSK simulations to lower-limb cycling were included; inverse kinematics-only was excluded. No protocol was registered, and no formal risk-of-bias assessment was conducted, as there were no intervention effects and no quantitative synthesis. Twenty-eight studies met the inclusion criteria. Most of them investigated bicycle–rider configuration, neuromuscular coordination, or electrical stimulation control, with participant cohorts overwhelmingly composed of young men and minimal female representation (272 total). Model reporting was often incomplete, with wide variation in anatomical scope, inconsistent descriptions of degrees of freedom, and limited sharing of models or code. Use of experimental data was uneven across studies: while all incorporated kinematic measurements, only two-thirds included kinetic data, and only one study reported physiological measures. Model validation was generally based on literature values. Seventy-eight per cent of studies used optimisation, mainly with effort-based cost functions, and parameter variations were exploratory rather than systematic. The evidence base is limited by small, predominantly male cohorts, inconsistent reporting standards, and limited physiological validation. These results consolidate current practices and highlight the need for more transparent and open reporting, sex-balanced and clinically diverse participant representation, stronger validation, and more rigorous sensitivity analysis to enhance reproducibility and practical relevance. This review was funded by AGAUR (Spain), CAPES (Brazil) and FAP-DF (Brazil).

**Author summary:** Cycling is widely used in sports training, rehabilitation, and clinical practice, and musculoskeletal simulations are increasingly used to study how muscles and joints work during cycling. These simulations allow us to estimate internal biomechanical variables that cannot be directly measured in experiments, such as muscle forces and joint loading. However, it is currently unclear how consistently these simulations are applied, validated, and reported across the literature. In this study, we systematically reviewed research published over the past 15 years that used musculoskeletal simulations to analyse lower-limb cycling. We identified 28 relevant studies and examined their modelling choices, experimental inputs, optimisation strategies, and validation approaches. We found substantial variability in model complexity, limited transparency in reporting, and a strong reliance on simplified literature-based validation methods. Most studies focused on narrow participant groups and explored modelling parameters in an *ad hoc* rather than a systematic way. Our findings highlight important gaps in current practice and point to clear opportunities for improvement. We provide an overview of common approaches and their limitations, and outline key recommendations to enhance the transparency, reproducibility, and practical relevance of musculoskeletal simulations in cycling research.

## Introduction

Cycling is widely recognised as a versatile activity that supports rehabilitation, promotes cardiovascular health, and reduces the risk of chronic diseases [1]. At a societal level, cycling encourages active mobility, contributes to environmental sustainability, and fosters community well-being [2]. However, optimising these benefits requires a detailed understanding of the mechanical demands imposed on the musculoskeletal system, particularly to enhance performance, reduce injury risk, and inform the design of bicycles, ergometers, and assistive technologies. Understanding the biomechanics of cycling is therefore of substantial interest across clinical, athletic, and engineering domains.

Research in cycling biomechanics has expanded over the past decades, producing detailed descriptions of pedalling kinematics, joint coordination, and muscle recruitment [3–8]. Experimental studies using inverse kinematics, inverse dynamics, and electromyography (EMG) have shown that the distribution of mechanical power and joint moments across the hip, knee, and ankle varies systematically with cadence and power output [3], and that fatigue alters muscle activation timing and inter-joint coordination, leading to compensatory strategies that reduce mechanical efficiency [4]. Complementary work has examined how posture, environmental conditions, and bike configuration influence performance and injury risk [5, 6]. While these findings have advanced our understanding of cycling biomechanics, experimental measurements inherently focus on directly observable phenomena. As a result, they provide limited access to internal variables, such as muscle forces and joint loading, and offer only constrained means to explore alternative strategies or equipment configurations.

Computational musculoskeletal simulations are increasingly used to address these limitations, enabling the estimation of internal mechanical variables that are difficult or impossible to measure directly [9]. Additionally, they allow controlled “what-if” investigations into how changes in technique, equipment, or neuromuscular capacity may influence performance, injury risk, or rehabilitation outcomes. More advanced approaches, including predictive simulations that generate optimal or emergent cycling motions [10], offer further opportunities to test strategies beyond the scope of experimental protocols. Recent perspectives emphasise that the value of these simulations lies not only in predicting motion, but also in clarifying underlying mechanisms and supporting decision-making in sport and clinical practice [11].

Despite these developments, it remains unclear which areas of cycling have most benefited from musculoskeletal simulation, and how differences in modelling choices, cycling protocols, and participant characteristics shape the scope, validity, and generalisability of current findings. Previous reviews have primarily focused on experimental pedalling biomechanics [12–15], muscle activation patterns [12], and injury-related factors [16, 17]. Broader reviews of predictive modelling exist [18], but they span multiple movement domains and do not examine cycling-specific modelling and simulation practices. As a result, we currently lack a systematic, simulation-focused synthesis that clarifies how modelling frameworks are being used, what applications they support, and how methodological choices affect reproducibility and interpretation.

Taken together, existing research demonstrates that cycling is a biomechanically rich activity, with well-documented relationships between pedalling mechanics, joint coordination, muscle activation, and task demands such as cadence, power output, and fatigue [3, 4]. Experimental approaches have provided essential insight into observable movement patterns and coordination strategies [5, 6], while computational musculoskeletal simulations extend this understanding by enabling access to internal mechanical variables and systematic exploration of hypothetical scenarios [9]. However, the rapid growth of simulation-based cycling studies has occurred alongside substantial methodological diversity, spanning modelling frameworks, control strategies, experimental protocols, and participant populations [10, 18]. Without a structured synthesis of how these approaches are being applied and evaluated, it remains difficult to assess the current state of the field, compare findings across studies, and identify both robust practices and critical gaps [11].

To bridge these gaps, we conducted a PRISMA-guided systematic review focused specifically on musculoskeletal simulations of lower-limb cycling. We examine not only the applications of these simulations, but also the modelling and analytical decisions that underpin them. This leads to two research questions:

1. *Q1: What are the main applications of musculoskeletal cycling simulations, and how have the study design, participant profiles, and cycling protocols influenced their scope and generalisability?*
2. *Q2: What modelling frameworks and analytical strategies have been adopted in musculoskeletal cycling simulations, and what do these choices reveal about current practices in terms of complexity, validation, and reproducibility?*

To address these questions, we systematically synthesise the existing literature on musculoskeletal cycling simulations. Specifically, we analyse how study design, participant characteristics, and cycling protocols shape the scope and generalisability of reported findings, as well as the modelling frameworks, control strategies, and validation approaches adopted across studies. By jointly considering the purposes for which these simulations are used and their implementation, this review provides a structured overview of current practices in simulation-based cycling biomechanics. The resulting synthesis highlights methodological trends, sources of variability, and gaps in validation and reproducibility, offering guidance for the design, interpretation, and future development of musculoskeletal cycling simulations in sport, clinical, and rehabilitation contexts.

## Materials and methods

This systematic review followed the PRISMA guidelines [19]. Fig 1 summarises the identification, screening, and selection process. The research question was structured using the Population/Problem, Concept, and Context (PCC) framework, adapted for computational modelling: here, the “Population” refers to the biomechanical problem addressed by simulations rather than specific human populations.

**Fig 1.**
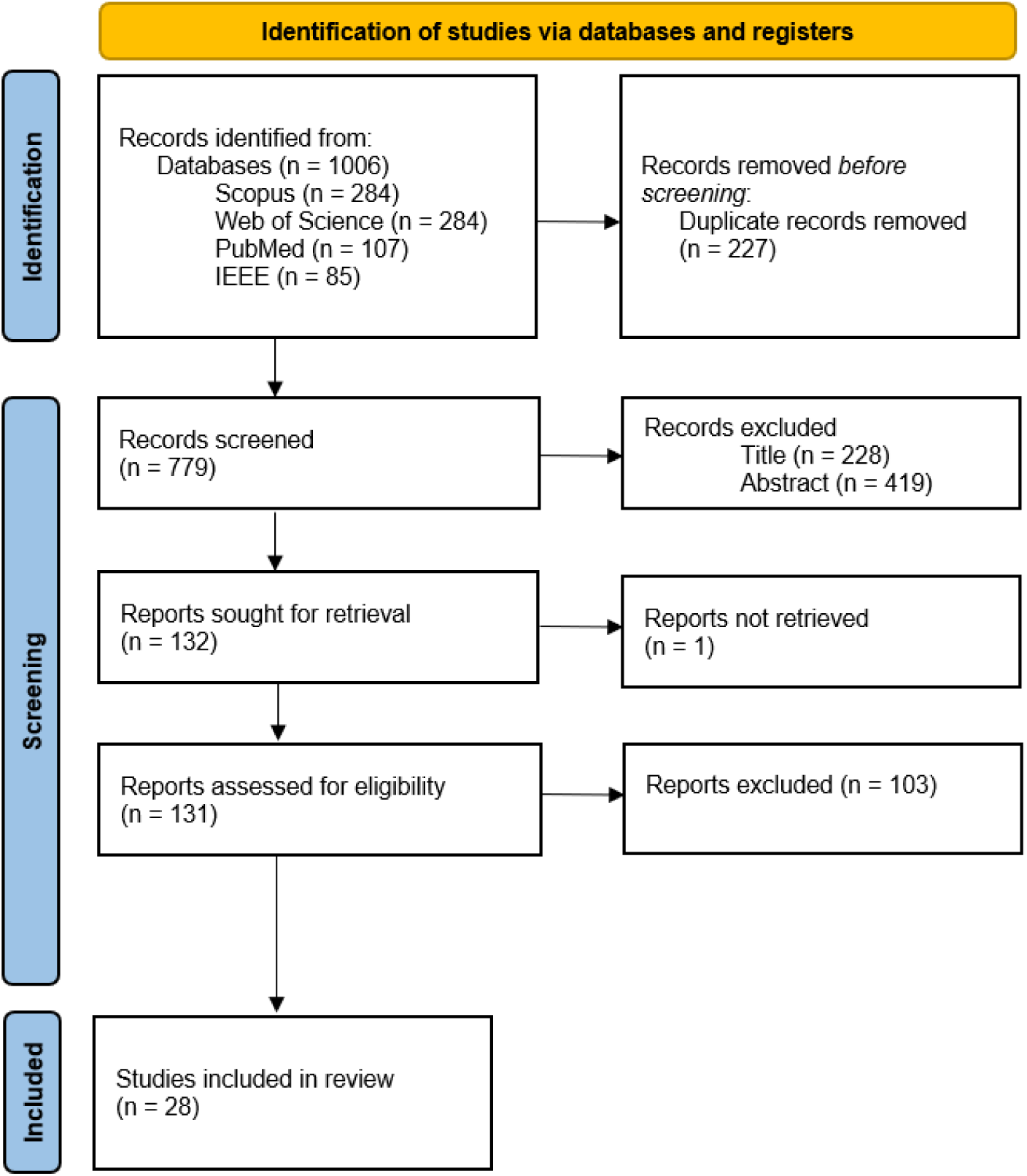
PRISMA flow diagram for study identification, screening, and inclusion.

### Eligibility criteria

Eligibility was determined using predefined inclusion and exclusion criteria (Table 1). We included peer-reviewed English-language journal articles that applied musculoskeletal simulations to analyse lower-limb internal biomechanics during cycling for performance, rehabilitation, or injury-related purposes. Articles were excluded if they lacked a musculoskeletal model, focused on non-cycling tasks, or reported only descriptive or inverse kinematics-only analyses.

**Table 1.** Inclusion and exclusion criteria for study selection.

| Inclusion criteria | Exclusion criteria |
| --- | --- |
| **Publication criteria** |  |
| Peer-reviewed journal articles available in English. | Non-peer-reviewed work (e.g., theses, opinions, reviews); unavailable full text; duplicates. |
| **Relevance to topic** |  |
| Cycling-focused lower-limb biomechanics. | Not related to cycling biomechanics; focused on non-cycling tasks or upper-body cycling. |
| Performance, rehabilitation, or injury-related applications. | Not addressing performance, rehabilitation, or injury. |
| **Methodological requirements** |  |
| Musculoskeletal simulation used. | No musculoskeletal model; inverse kinematics-only or EMG-only analyses. |
| Simulation approaches such as static optimisation, forward dynamics, or tracking. | No simulation of internal biomechanics. |
| **Biomechanical focus** |  |
| Internal mechanical variables (e.g., muscle forces, joint loading). | No internal biomechanical outcomes. |

### Information sources

Searches were conducted on Scopus, PubMed, IEEE Xplore, and Web of Science on 1 August 2024. No grey literature, organisational websites, or additional registers were included.

### Search strategy

The search strategy combined terms related to cycling, biomechanics, musculoskeletal modelling, simulation techniques, and relevant biomechanical variables (Table 2). Boolean operators and proximity constraints (e.g., W/15) were used to improve specificity. Searches were conducted on titles and abstracts of peer-reviewed publications published between January 2010 and July 2024 and restricted to English-language articles.

**Table 2.**
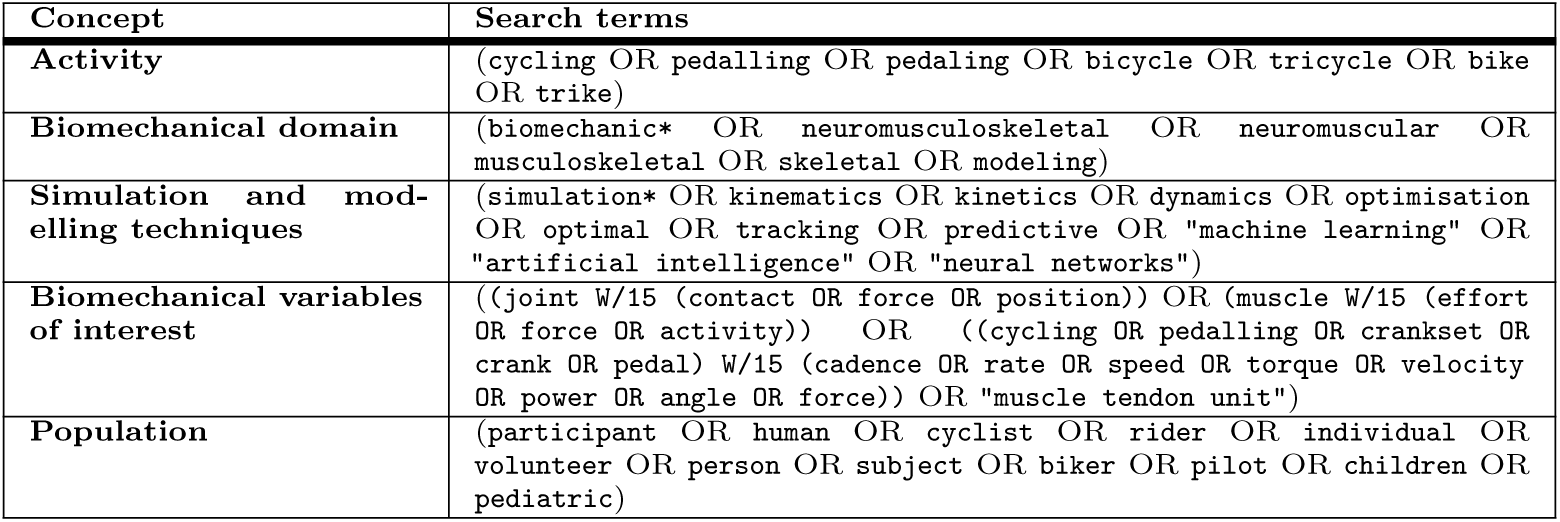
Search terms grouped by conceptual category used to construct the database queries. Proximity operator W/15 requires terms to appear within 15 words of each other.

| Concept | Search terms |
| --- | --- |
| Activity | (cycling OR pedalling OR pedaling OR bicycle OR tricycle OR bike OR trike) |
| Biomechanical domain | (biomechanic* OR neuromusculoskeletal OR neuromuscular OR musculoskeletal OR skeletal OR modeling) |
| Simulation and modelling techniques | (simulation* OR kinematics OR kinetics OR dynamics OR optimisation OR optimal OR tracking OR predictive OR "machine learning" OR "artificial intelligence" OR "neural networks") |
| Biomechanical variables of interest | ((joint W/15 (contact OR force OR position)) OR (muscle W/15 (effort OR force OR activity)) OR ((cycling OR pedalling OR crankset OR crank OR pedal) W/15 (cadence OR rate OR speed OR torque OR velocity OR power OR angle OR force)) OR "muscle tendon unit") |
| Population | (participant OR human OR cyclist OR rider OR individual OR volunteer OR person OR subject OR biker OR pilot OR children OR pediatric) |

### Selection process

All records underwent title, abstract, and full-text screening. Reviewers independently screened an equal portion of titles and abstracts; the first author double-checked exclusions. All authors independently reviewed full texts, and disagreements were resolved through discussion. All screening steps were managed via shared Google Sheets.

### Data collection process

Data were independently extracted by all reviewers using a review-specific, standardised Google Sheets template developed by the authors. Extracted entries were compared and consolidated through consensus. No automated extraction tools were used.

### Data items

This review extracted variables describing the aims, methodological choices, computational approaches, and analytical frameworks used across studies.

#### Study aims, context, and participants’ characteristics

Practical application domains were categorised as rehabilitation, sports performance, or injury-related. The purpose of each simulation was identified (e.g., bicycle-rider configuration analysis, neuromuscular function and muscle coordination, functional electrical stimulation (FES) and rehabilitation, and computational model development and validation).

Cycling protocols were described using cadence and power output, which were the most consistently reported variables. Studies were additionally categorised by bicycle or ergometer type (seated ergometer, upright ergometer, recumbent and upright road bike, Fig 2).

**Fig 2.**
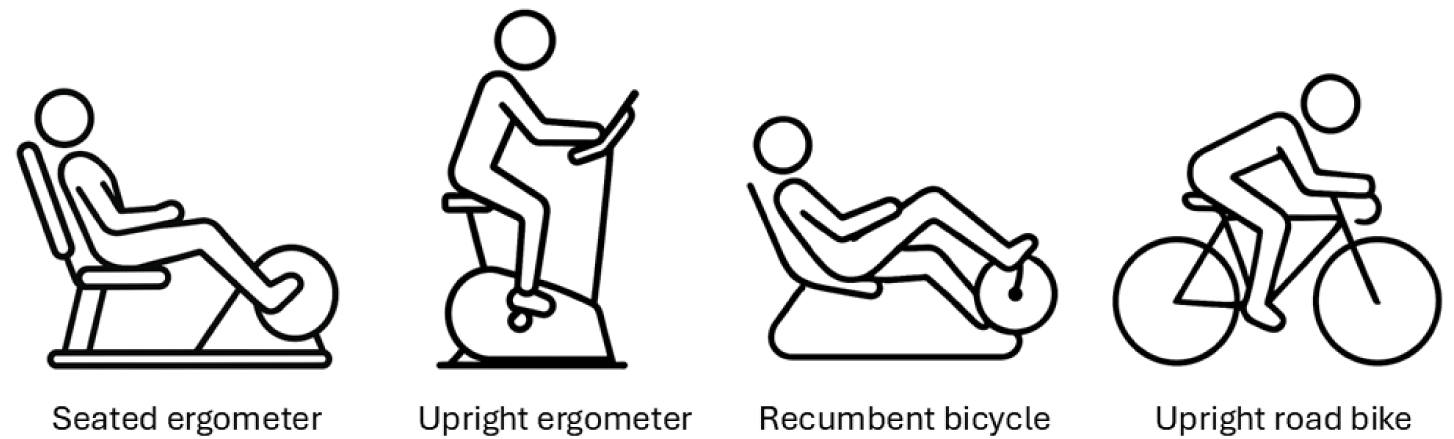
Schematic representation of the bicycle and ergometer types used to categorise cycling protocols across studies. These configurations differ in rider posture, support, and crank–seat geometry, which may influence biomechanical demands, muscle activation patterns, and power production.

Participant characteristics included demographic (age, sex, number), anthropometric (body mass, height, BMI), cycling ability (recreational, trained, professional), and clinical (e.g., patients with spinal cord injury or knee osteoarthritis) attributes.

#### Methodological frameworks

Models were classified by anatomical scope (single-leg, two-leg, full-body without arms, full-body with arms), dimensionality (2D/3D), degrees of freedom (DOF), and number of muscle–tendon units.

Software platforms were recorded (custom, OpenSim, AnyBody), along with additional optimisation tools (e.g., Moco, FAST, SNOPT). Code availability was verified using the provided links or repositories.

The experimental data used for comparison were grouped into kinematic, kinetic, neuromuscular, physiological, or loading-related categories.

Computational simulation methods were categorised as inverse kinematics (when extended to musculoskeletal analysis; otherwise, the paper was excluded), static optimisation, computed muscle control, forward dynamics, optimal control, or hybrid approaches.

Cost function terms were classified into the following categories: minimisation of muscle activity or effort, minimisation of metabolic energy, tracking of pedal kinematics or forces, maximisation of power or joint work, minimisation of joint reaction forces, tracking of joint moments, joint angles, or EMG signals, and an “other” category for cost function terms that did not fit within these classifications.

Sensitivity and perturbation analyses were classified into the following categories: geometric or equipment-related, workload or task-related, musculoskeletal or morphological, joint or kinematic, and control or cost function-related perturbations. An “other” category was used for perturbations that did not fit within these predefined classes. When studies included multiple perturbation types, a multi-label classification approach was applied.

### Reporting transparency and PRISMA limitations

This review followed PRISMA 2020 guidelines [19]; however, several checklist items do not apply to our study design or were not fully addressed. First, no protocol was prospectively registered, and, therefore, no registration number is available. Although Table 1 defines the eligibility criteria, the full search strings executed in each database are not included in the manuscript and are instead provided in the Supplementary material S1 for completeness.

Because the objective of this review was to map methodological practices rather than synthesise intervention effects, we did not perform a formal risk-of-bias assessment. Likewise, no effect measures, meta-analytic combinations, or certainty assessments were planned or conducted. Narrative synthesis procedures were applied, but we did not explicitly describe the criteria guiding study grouping or thematic organisation. An assessment of reporting bias was also not undertaken, as this PRISMA item is relevant primarily to quantitative effect synthesis and meta-analysis, which were outside the scope of this review.

Together, these clarifications outline the scope of PRISMA requirements relevant to this methodological review and specify the items intentionally not undertaken.

## Results

The initial search across Scopus (284 records), Web of Science (530), PubMed (107), and IEEE Xplore (85) yielded a total of 1,006 records. After removing duplicates, 779 unique records remained and underwent title screening, which excluded 228 articles. The remaining 551 records proceeded to abstract screening, resulting in the exclusion of an additional 419 records (e.g., [20], which did not implement a musculoskeletal muscle-driven model). A total of 132 articles were then sought for full-text retrieval; one report was inaccessible. Of the 131 full-text articles assessed for eligibility, 103 were excluded according to predefined criteria. Ultimately, 28 studies met all inclusion criteria and were incorporated into the final review.

### Study aims, context, and participant characteristics

#### Application domains and purpose of musculoskeletal simulation

Across the included studies, practical applications often overlapped: 53.6% (15/28) addressed rehabilitation, 64.3% (15/28) targeted sports performance, and 32.1% (9/28) examined injury-related questions (Fig 3 and 4(A)).

**Fig 3.**
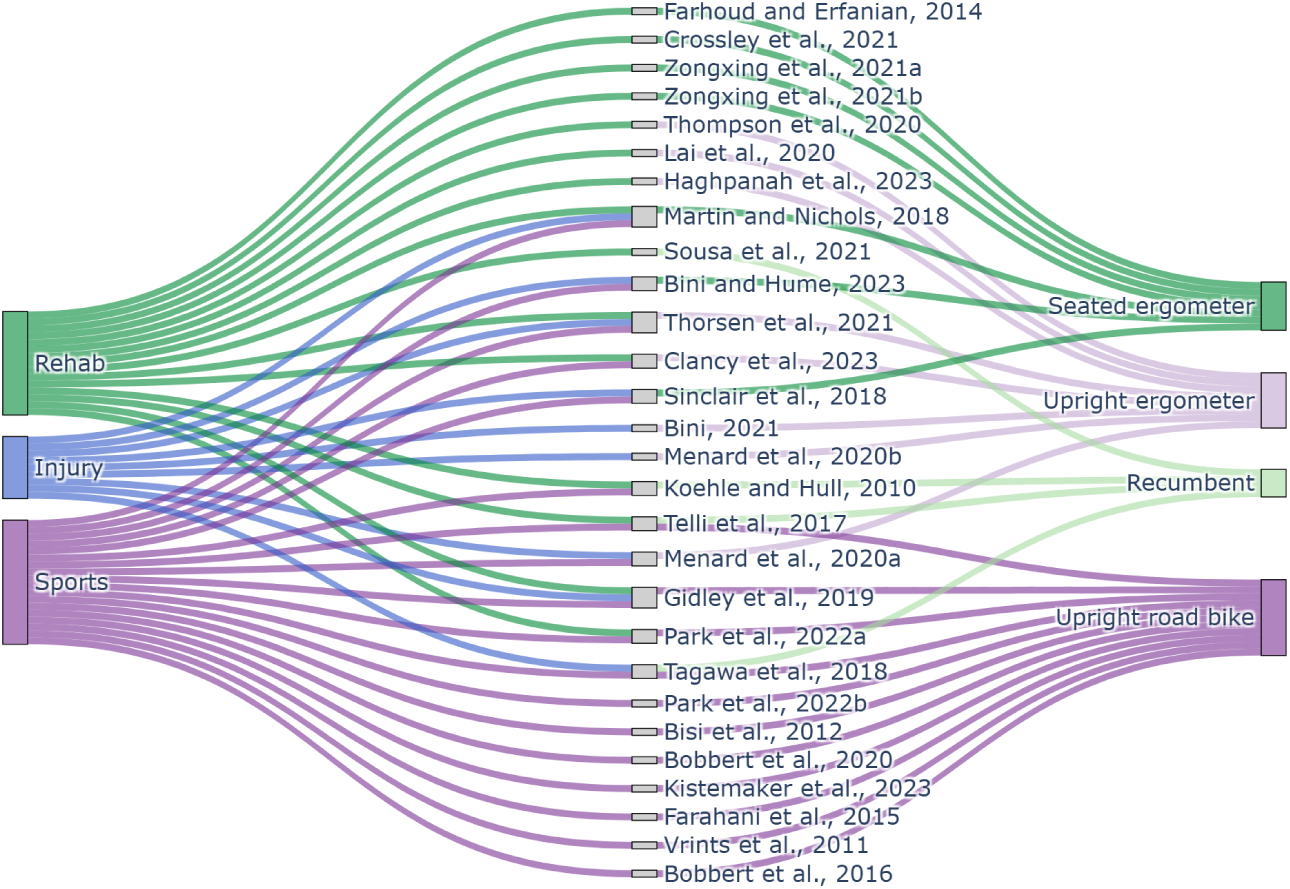
Relationship between study application domain, individual studies, and cycling equipment used in musculoskeletal simulation research. Sankey diagram showing how rehabilitation, injury, and sports-focused studies map onto specific publications and cycling configurations, including seated ergometers, upright ergometers, recumbent bikes, and upright road bikes. Flow widths indicate the number of studies associated with each category.

Four overarching purposes for musculoskeletal simulations in cycling emerged. **First**, a large group of studies focused on bicycle–rider configuration and equipment optimisation, using simulations to evaluate how systematic variations in setup parameters influence biomechanical outcomes. These studies examined changes in seat-tube angle [21], saddle position [22–25], crank length [26], Q-factor [27], chainring shape [28], and recumbent versus upright posture [29]. Across these configurations, simulations were used to assess outcomes such as joint loading and joint reaction forces [30], radial pedal force [31], force–velocity relationships [32], and injury-related metrics including iliotibial band syndrome risk [33]. **Second**, a cluster of studies examined neuromuscular function and muscle coordination, using simulations to study muscle activation strategies [34], contractile behaviour [35], biarticular contributions [36], or the influence of muscle morphology on performance [37]. **Third**, some investigations targeted functional electrical stimulation (FES) and rehabilitation control, which used simulations to optimise stimulation strategies [38], evaluate assistive devices such as orthoses [39], examine joint loading in clinical populations [40], or design hybrid training for altered gravity [41]. **Finally**, a small group focused on computational model development and validation, including the creation or validation of direct-collocation frameworks [42–44], comparisons of knee joint models [45], predictions of crank torque or pedal motion [46], and the provision of open datasets and reproducible modelling pipelines [47, 48].

**Fig 4.**
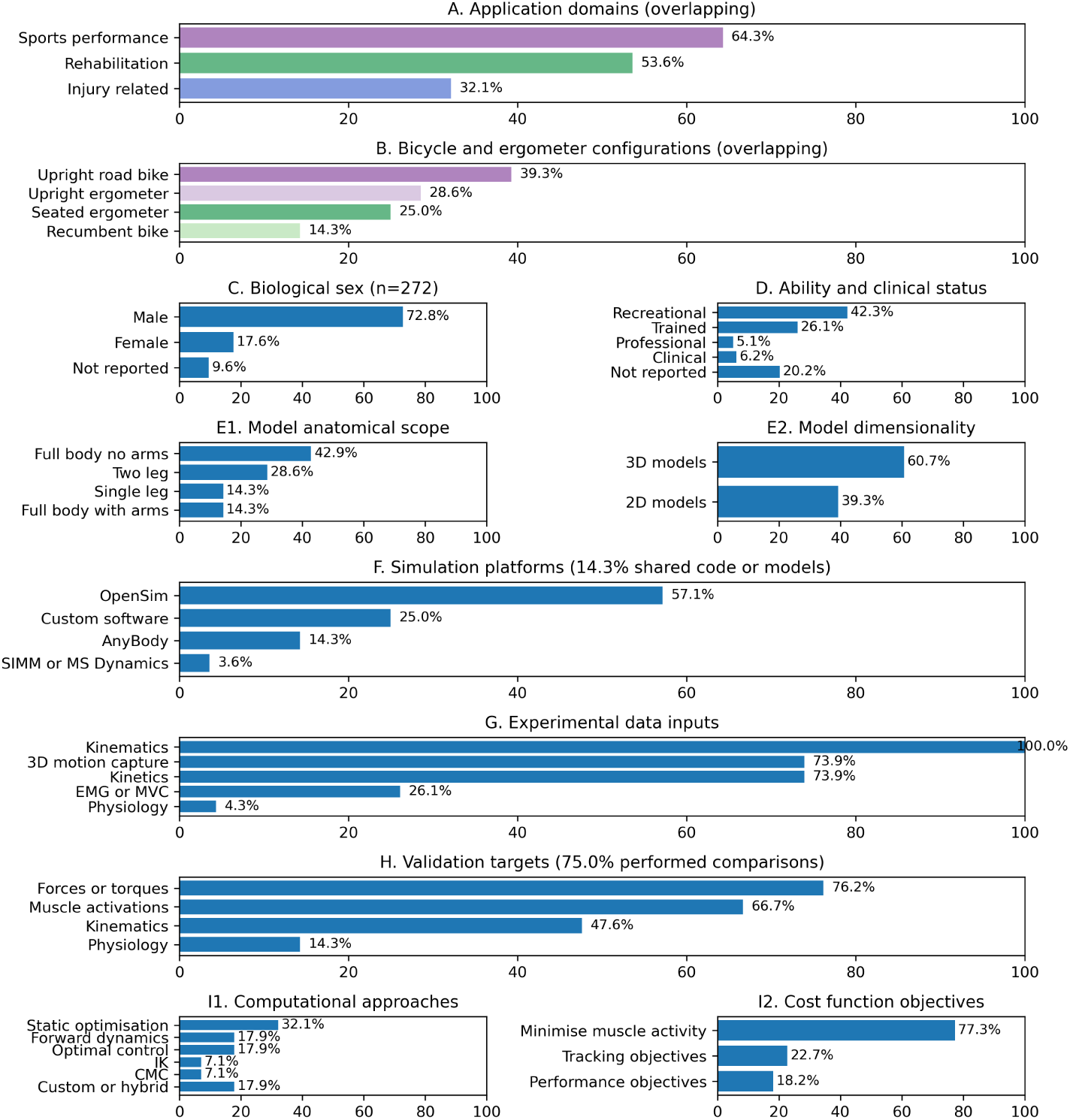
Overview of study characteristics and modelling practices in musculoskeletal cycling simulations. Summary of application domains and equipment configurations (A–B), participant characteristics (C–D), model scope and dimensionality (E1–E2), simulation platforms and code availability (F), experimental data inputs (G), validation targets (H), computational approaches (I1), and cost function objectives (I2) across the included studies. Application domains and bicycle/ergometer configurations could overlap within studies (A–B). Participant distributions (C–D) are reported as percentages of the total number of participants (n = 272). Model characteristics, simulation platforms, and computational approaches (E–F, I1) are reported as percentages of studies. Experimental data inputs (G) are reported relative to studies that collected experimental data, while validation targets (H) are reported relative to studies that performed any comparison between simulations and experimental or literature data; the proportion of studies performing such comparisons is indicated in the panel title. cost function objectives (I2) are reported relative to studies with an explicitly defined optimisation objective, excluding studies without a cost function.

#### Cycling protocol and bicycle type

Cycling protocols showed trends across application domains. Rehabilitation and FES studies predominantly used low cadences (30–60 rpm) and low power outputs (15–45 W) (e.g., [38]), aligning with reduced physical capacity or safety considerations. Sports and injury-focused studies more often used moderate cadences (60–90 rpm) and moderate power levels (60–100 W) (e.g., [30]), while trained or professional cyclists were the only groups to reach *<90 rpm and <100* W (e.g., [48]). Simulation-only studies consistently operated at free cadence and power (e.g., [31, 41]).

Equipment selection also varied systematically across application domains (Fig 3). Overall, 39.3% (11/28) of studies used upright road bikes, 14.3% (4/28) used recumbent bikes, 28.6% (8/28) used upright ergometers and 25.0% (7/28) used seated ergometers (Fig 4(B)). These distributions largely mirrored the intended application. Rehabilitation and FES-oriented studies predominantly relied on seated ergometers and recumbent configurations, reflecting safety, stability, and accessibility considerations. In contrast, sports-focused studies most frequently employed upright road bikes, consistent with performance-oriented objectives and ecological validity. Injury-related studies spanned both ergometer-based and upright configurations, reflecting their intermediate positioning between controlled assessment and sport-specific relevance.

#### Participant characteristics

The studies represented data from 272 participants (Table 3), though not all were unique, as some datasets were reused from previous publications ([30, 35, 36, 42, 43, 45, 47]). Biological sex was reported for 246 participants, comprising 198 males (72.8%) and 48 females (17.6%), while sex was not described for 26 participants (9.6%) (Fig 4(C)). Cycling ability or clinical condition was reported for 217 participants: 115 recreational (42.3%), 71 trained (26.1%), and 14 professional cyclists (5.1%) (Fig 4(D)). Clinical samples included 4 participants with spinal cord injury (1.5%) and 13 with knee osteoarthritis (4.8%). Cycling ability was not described for 55 participants (20.2%).

**Table 3.** Participant characteristics reported. Summary of participant characteristics reported in each included study, including sample size, sex distribution, participant ability level, and clinical status.

| Reference | N Participants | Male | Female | Recreational | Trained | Professional | SCI | OA |
| --- | --- | --- | --- | --- | --- | --- | --- | --- |
| Park et al., 2022a [42] | 15 | 11 | 4 | 15 | 0 | 0 | 0 | 0 |
| Park et al., 2022b [34] | 15 | 11 | 4 | 15 | 0 | 0 | 0 | 0 |
| Bisi et al., 2012 [21] | 10 | 10 | 0 | NA | NA | NA | NA | NA |
| Bobbert et al., 2020 [37] | 0 | 0 | 0 | 0 | 0 | 0 | 0 | 0 |
| Farhoud and Erfanian, 2014 [38] | 3 | 3 | 0 | 0 | 0 | 0 | 3 | 0 |
| Menard et al., 2020a [33] | 10 | NA | NA | 0 | 10 | 0 | 0 | 0 |
| Thorsen et al., 2021 [27] | 16 | 8 | 8 | 16 | 0 | 0 | 0 | 0 |
| Bini, 2021 [23] | 10 | 6 | 4 | 10 | 0 | 0 | 0 | 0 |
| Menard et al., 2020b [22] | 10 | 10 | 0 | 0 | 10 | 0 | 0 | 0 |
| Crossley et al., 2021 [40] | 11 | NA | NA | 11 | 0 | 0 | 0 | 0 |
| Kistemaker et al., 2023 [31] | 0 | 0 | 0 | 0 | 0 | 0 | 0 | 0 |
| Thompson et al., 2020 [30] | 24 | 12 | 12 | 11 | 0 | 0 | 0 | 13 |
| Lai et al., 2020 [35] | 20 | 10 | 10 | 0 | 20 | 0 | 0 | 0 |
| Clancy et al., 2023 [47] | 16 | 11 | 5 | NA | NA | NA | NA | NA |
| Haghpanah et al., 2023 [44] | 5 | NA | NA | NA | NA | NA | NA | NA |
| Sousa et al., 2021 [39] | 1 | 1 | 0 | 0 | 0 | 0 | 1 | 0 |
| Gidley et al., 2019 [43] | 24 | 24 | 0 | NA | NA | NA | NA | NA |
| Farahani et al., 2015 [46] | 8 | 8 | 0 | 0 | 8 | 0 | 0 | 0 |
| Telli et al., 2017 [29] | 4 | 4 | 0 | 4 | 0 | 0 | 0 | 0 |
| Bini and Hume, 2023 [48] | 17 | 17 | 0 | 3 | 0 | 14 | 0 | 0 |
| Martin and Nichols, 2018 [36] | 13 | 12 | 1 | 0 | 13 | 0 | 0 | 0 |
| Tagawa et al., 2018 [41] | 0 | 0 | 0 | 0 | 0 | 0 | 0 | 0 |
| Zongxing et al., 2021a [26] | 0 | 0 | 0 | 0 | 0 | 0 | 0 | 0 |
| Koehle and Hull, 2010 [45] | 15 | 15 | 0 | 15 | 0 | 0 | 0 | 0 |
| Vrints et al., 2011 [25] | 10 | 10 | 0 | 0 | 10 | 0 | 0 | 0 |
| Zongxing et al., 2021b [24] | 0 | 0 | 0 | 0 | 0 | 0 | 0 | 0 |
| Sinclair et al., 2018 [28] | 15 | 15 | 0 | 15 | 0 | 0 | 0 | 0 |
| Bobbert et al., 2016 [32] | 0 | 0 | 0 | 0 | 0 | 0 | 0 | 0 |
| **Total** | **272** | **198** | **48** | **115** | **71** | **14** | **4** | **13** |

*Note:* Values reported as ‘0’ indicate explicitly reported absence, whereas ‘NA’ denotes information not reported.

Across studies, participants were predominantly young adults, typically aged 24–30 years. Only a small number included middle-aged adults or wider age ranges (e.g., [30]). Anthropometric characteristics were comparatively homogeneous, with most studies reporting heights between 1.70 and 1.77 m and body masses between 65 and 77 kg, aside from clinical groups [30, 39, 45, 47], which tended to have higher and more variable body mass.

### Methodological frameworks

#### Type of model and dimensionality

Musculoskeletal models showed substantial variation in anatomical scope and dimensionality (Table 4). Four studies (14.3%) used single-leg models (1L), eight (28.6%) used two-leg models (2L), four (14.3%) employed full-body models including the arms (FB), and twelve (42.9%) used full-body models without arms (FBna) (Fig 4(E1)). Regarding dimensionality, 39.3% of models were 2D, and 60.7% were 3D (Fig 4(E2)). The dimensionality was not always explicitly stated and was often inferred from out-of-plane kinematics, such as hip abduction–adduction.

**Table 4.**
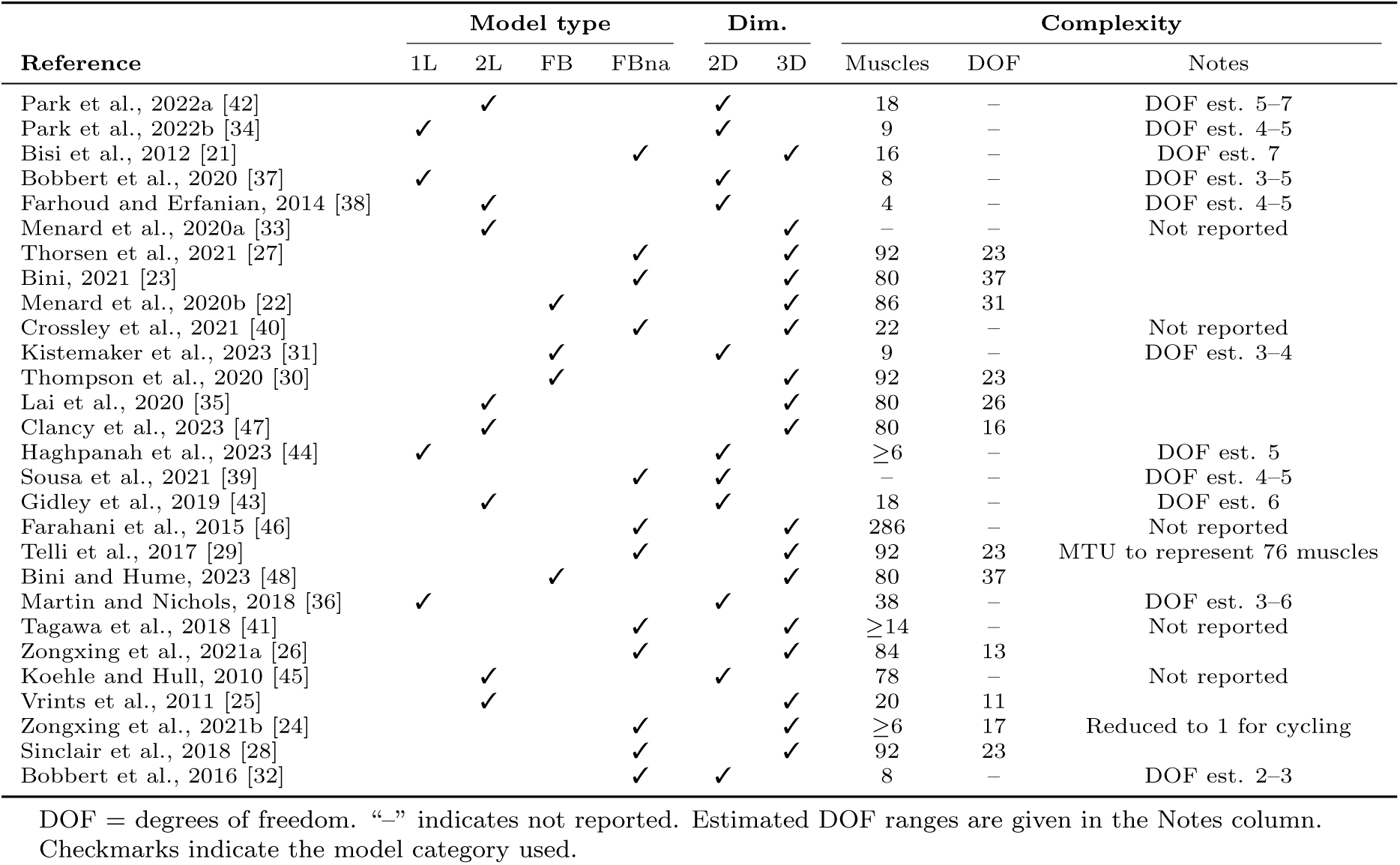
Overview of modelling characteristics. Reported variables include model configuration, dimensionality, number of muscles, and number of degrees of freedom (DOF). Model type abbreviations: 1L – single-leg model; 2L – bilateral two-leg model; FB – full-body model; FBna – full-body model with non-actuated segments. Dimensionality refers to whether the model is two-dimensional (2D) or three-dimensional (3D). Missing information is indicated as “Not reported.”

#### Number of muscles and degrees of freedom

Muscle–tendon unit (MTUs) definitions varied widely (Table 4), ranging from 4 (e.g., [38]) to 286 (e.g., [46]) muscle–tendon units, and many studies did not provide a full list of included muscles. Reported or inferred degrees of freedom ranged from 1 (e.g., [26]) to 37 (e.g., [48]), yet approximately half of the studies did not specify the degrees of freedom (DOFs) used. Ambiguities were common regarding whether DOFs referred to the base model, the adapted cycling configuration, or the effective closed-chain DOFs. These inconsistencies limited direct comparison of model complexity across studies.

#### Software platform used and code and model availability

OpenSim was the most commonly used platform (57.1%, 16/28), followed by custom software implementations (25.0%, 7/28), AnyBody (14.3%, 4/28) and SIMM and Musculoskeletal Dynamics (3.6%, 1/28) (Fig 4(F) and Table 5). Despite the predominance of open-source software, only four studies (14.3%, [39, 42, 43, 47]) made model or code resources publicly available. Availability was determined based on explicit statements in the manuscript, supplementary or complementary materials, and URLs provided within the article. The vast majority did not provide accessible implementations, even when availability was suggested.

**Table 5.** Software platforms used across included studies. References are grouped by computational platform. Categories are not mutually exclusive.

| Software | References |
| --- | --- |
| OpenSim | [42] [34] [33] [27] [23] [22] [40] [30] [35] [47] [39] [29] [48] [36] [25] [28] |
| Custom | [21] [37] [38] [31] [44] [43] [32] |
| AnyBody | [46] [41] [26] [24] |
| SIMM/MSD | [45] |

#### Experimental data collection

Among the 28 studies, 15 (53.6%) collected new experimental data, eight (28.6%) used previously collected datasets, and five (17.9%) did not include experimental data (Table 6). Of the 23 studies using experimental data, all reported kinematic measurements (100%, 23/23), ranging from simple IMU-based crank angle recordings to full 3D motion capture (73.9%, 17/23). Kinetic data were reported in 17 studies (73.9%), all of which measured crank or pedal forces; only one study also included seat forces. Neuromuscular activation data (EMG or Maximum Voluntary Contraction (MVC)) were collected in 6 studies (26.1%). Physiological or metabolic measurements were rare, appearing in only one study (4.3%, [21]) (Fig 4(G)).

Among the 21 studies (75.0%) that compared simulated outcomes with experimental or literature data, 42.9% (9/21) relied partly on literature data (Table 6). Simulated kinematics were compared against experimental measurements in 47.6% of studies (10/21), forces or torques in 76.2% (16/21), and muscle activations in 66.7% (14/21). Physiological comparisons were performed in 14.3% (3/21) (Fig 4(H)). No study directly validated simulated loading conditions against experimentally measured resistance profiles.

#### Computational simulation

Across the 28 studies, a wide range of computational approaches was used (Table 7). Static optimisation (SO) was the most common method (32.1%, 9/28), followed by forward dynamics (FD) and optimal control (OC), each implemented in 17.9% of studies (5/28) (Fig 4(I1)). Among the forward dynamics studies, some incorporated genetic algorithm (GA)–based optimisation within the simulation [32, 37, 44]. Optimal control approaches employed a variety of solvers, including IPOPT [34, 42, 47], SNOPT [31], and simulated annealing (SA) [43], and were used in both tracking and predictive simulations. Inverse kinematics (IK, [25, 33]) and computed muscle control (CMC, [28, 35]) were less frequent, each appearing in 7.1% of studies. Five studies (17.9%) used custom or hybrid computational frameworks that did not fit within these standard categories.

**Table 6.**
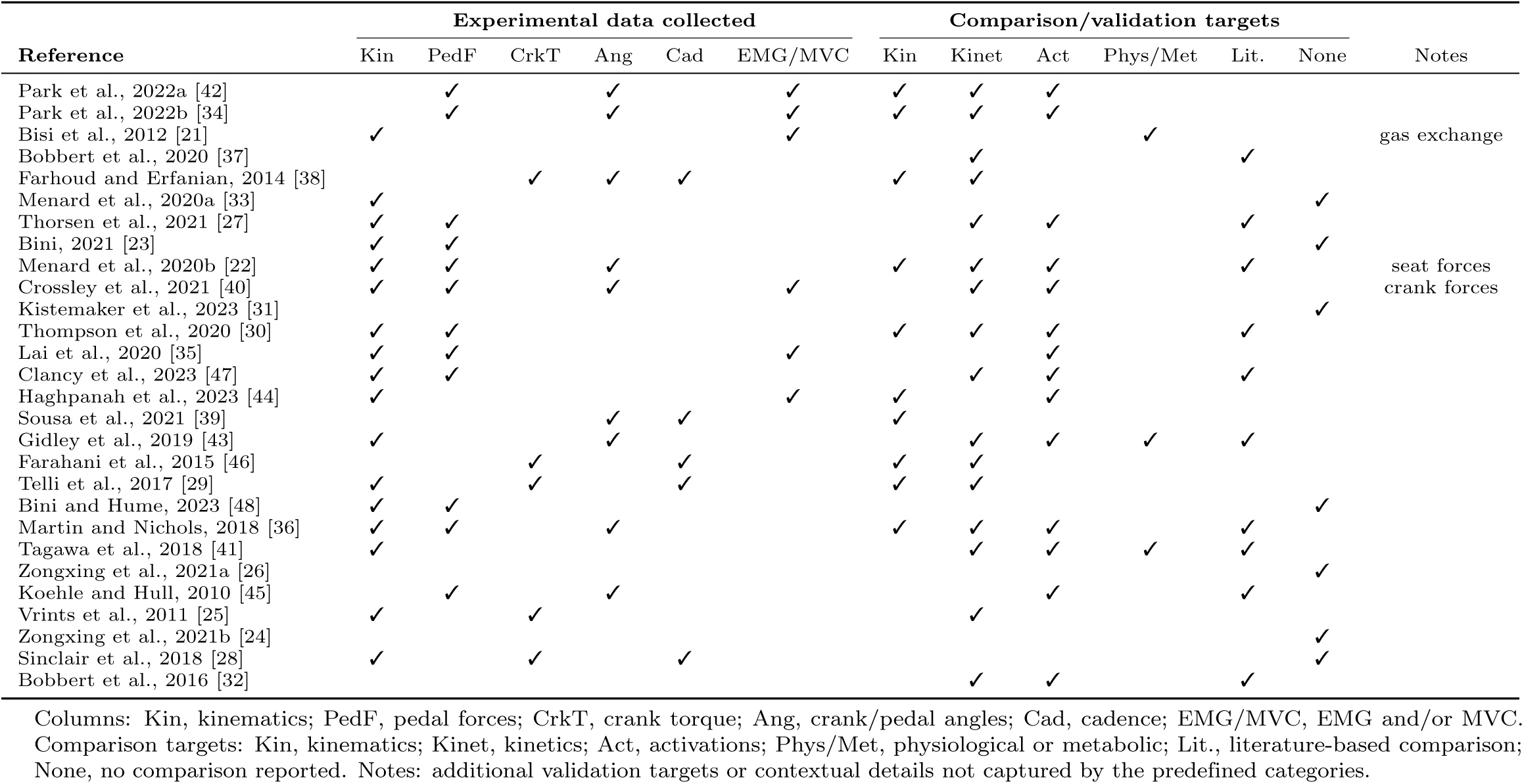
Overview of experimental data collected. Summary of the types of measurements acquired in the original experiments and the variables used for evaluation or validation of the musculoskeletal models. EMG denotes electromyography signals. Kinematics refers to full-body motion data, typically including joint angles and segment trajectories. Activations correspond to neuromuscular activations. In the comparison/validation section, “Lit.” indicates that model outputs were compared with data reported in previously published studies, rather than with experimental data collected within the same article.

**Table 7.** Simulation control methods and cost functions. Checkmarks indicate categories reported in each study.

| Reference | Control |  |  |  |  |  | Cost |  |  |  | Notes |
| --- | --- | --- | --- | --- | --- | --- | --- | --- | --- | --- | --- |
|  | DC | FD | SO | IK | CMC | EMG | minAct | Track | maxPower | EMGmatch |  |
| Park et al., 2022a [42] | ✓ |  |  |  |  |  | ✓ | ✓ |  |  | Track pedal forces and pedal angles |
| Park et al., 2022b [34] | ✓ |  |  |  |  |  | ✓ | ✓ |  |  | Track pedal forces and pedal angles |
| Bisi et al., 2012 [21] |  |  |  |  |  | ✓ |  |  |  |  |  |
| Bobbert et al., 2020 [37] |  | ✓ |  |  |  |  |  |  | ✓ |  | GA; Non-periodic penalty |
| Farhoud and Erfanian, 2014 [38] |  | ✓ |  |  |  |  |  |  |  |  | Fuzzy; Sliding mode |
| Menard et al., 2020a [33] |  |  |  | ✓ |  |  |  |  |  |  |  |
| Thorsen et al., 2021 [27] |  |  | ✓ |  |  |  | ✓ |  |  |  |  |
| Bini, 2021 [23] |  |  | ✓ |  |  |  | ✓ |  |  |  |  |
| Menard et al., 2020b [22] |  |  | ✓ |  |  |  | ✓ |  |  |  |  |
| Crossley et al., 2021 [40] |  |  |  |  |  | ✓ | ✓ | ✓ |  | ✓ | Track joint moments |
| Kistemaker et al., 2023 [31] | ✓ | ✓ |  |  |  |  |  |  | ✓ |  |  |
| Thompson et al., 2020 [30] |  |  | ✓ |  |  |  | ✓ |  |  |  |  |
| Lai et al., 2020 [35] |  |  |  |  | ✓ |  | ✓ |  |  |  |  |
| Clancy et al., 2023 [47] | ✓ |  |  |  |  |  | ✓ |  |  |  | Tibiofemoral JRF |
| Haghpanah et al., 2023 [44] |  | ✓ |  |  |  |  |  | ✓ |  |  | GA; CPG; Track trajectories |
| Sousa et al., 2021 [39] |  | ✓ |  |  |  |  |  |  |  |  |  |
| Gidley et al., 2019 [43] | ✓ |  |  |  |  |  | ✓ | ✓ |  |  | SA; Track kinematics; Energy; Stress; Smoothness |
| Farahani et al., 2015 [46] |  |  |  |  |  |  | ✓ |  |  |  | Inverse-inverse dynamics |
| Telli et al., 2017 [29] |  |  |  |  |  |  |  |  |  |  | MTU simulation |
| Bini and Hume, 2023 [48] |  |  | ✓ |  |  |  | ✓ |  |  |  |  |
| Martin and Nichols, 2018 [36] |  |  |  |  |  |  |  |  | ✓ |  | Work-loop optimisation |
| Tagawa et al., 2018 [41] |  |  | ✓ |  |  |  | ✓ |  |  |  |  |
| Zongxing et al., 2021a [26] |  |  | ✓ |  |  |  | ✓ |  |  |  |  |
| Koehle and Hull, 2010 [45] |  |  | ✓ |  |  |  | ✓ |  |  |  |  |
| Vrints et al., 2011 [25] |  |  |  | ✓ |  |  |  |  |  |  |  |
| Zongxing et al., 2021b [24] |  |  | ✓ |  |  |  | ✓ |  |  |  | Fatigue-based |
| Sinclair et al., 2018 [28] |  |  |  |  | ✓ |  | ✓ |  |  |  |  |
| Bobbert et al., 2016 [32] |  | ✓ |  |  |  |  |  |  | ✓ |  | GA |

Abbreviations: DC = direct collocation optimal control; FD = forward dynamics; SO = static optimisation; IK = inverse kinematics; CMC = computed muscle control; EMG = EMG driven method; GA = Genetic algorithm; SA = Simulated annealing; JRF = Joint reaction forces.
Cost terms: minAct = minimise muscle activation effort; Track = tracking term; maxPower = maximise power; EMGmatch = match experimental EMG.
Notes include additional objectives, penalties, or modifiers not represented as separate columns.

#### cost functions

Of the 28 studies reviewed, 22 incorporated an optimisation framework with an explicit cost function (Table 7) (Fig 4(I2)). Among these, minimising muscle activity or effort (minAct) was the most common objective (77.3%, 17/22). Several studies also included tracking-based terms (Track), such as tracking pedal angles or forces, joint moments, joint angles or EMG (22.7%, 5/22). Only one study explicitly incorporated experimental EMG matching as part of the optimisation objective ([40]), highlighting the limited use of data-driven neuromuscular validation within predictive frameworks. Objectives aimed at performance optimisation were less frequent, including maximising power or work output and maximising joint power or work (18.2%, 4/22) (maxPower). A small number of studies targeted physiological or structural outcomes, such as minimising joint reaction forces (minJRF) [47].

#### Sensitivity analysis

Of the 28 studies reviewed, 27 incorporated some form of parameter variation, while one study did not conduct any sensitivity or perturbation analysis [36] (Table 8). The most common category was geometric or equipment sensitivity, reported in nine studies (32.1%, 9/28), primarily examining how changes in bicycle configuration, such as saddle height or setback, crank length, seat-tube angle, pelvic orientation, or chainring geometry, affected biomechanical or muscle-level outcomes. Workload or task sensitivity appeared in six studies (21.4%, 6/28), typically by varying cadence, power output, or resistance to assess its influence on mechanical efficiency or muscular demand. Musculoskeletal or morphological sensitivity was also identified in six studies (21.4% 6/28), in which model properties, such as muscle morphology or joint structures, were systematically altered. Control or cost function sensitivity was observed in seven studies (25.0%, 7/28), representing the largest cluster among optimal control and predictive-simulation frameworks; these studies varied optimisation objectives, weighting factors, solver settings, or controller robustness. One study [34] fell into the other comparative category, as it employed a pre–post learning design rather than a formal perturbation analysis. Overall, although parameter variations were common, most studies did not explicitly frame these tests as formal sensitivity analyses; instead, they used them as exploratory tools to examine how geometric, physiological, or control modifications influenced simulation or experimental outcomes.

**Table 8.** Sensitivity categories identified across included studies. Categories are not mutually exclusive; some studies appear in multiple rows.

| Sensitivity category | References |
| --- | --- |
| Geometric / Equipment | [21] [33] [23] [22] [30] [26] [25] [24] [28] |
| Workload / Task | [40] [31] [35] [29] [48] [41] |
| Musculoskeletal / Morphological | [37] [39] [29] [41] [45] [32] |
| Control / Cost-function | [42] [38] [27] [47] [44] [43] [46] |
| Other / Comparative | [34] |
| No sensitivity analysis | [36] |

## Discussion

This systematic review synthesises the use of musculoskeletal simulations in cycling biomechanics over the past 15 years, revealing clear patterns in study aims, participant profiles, modelling strategies, and validation practices across 28 papers. The evidence shows a field that is technically diverse yet methodologically inconsistent, with substantial variation in model choice, parameter reporting and optimisation approaches, and persistent gaps in population diversity and reproducibility. These findings situate the current state of cycling simulation research, highlight how methodological decisions shape model outputs and interpretability, and identify when greater standardisation, transparency, and task-specific validation are needed to advance applications in performance, rehabilitation, and injury prevention.

### Study aims, context, and participant characteristics

Cycling simulations were used for a wide range of purposes, but most studies fell into four categories: equipment and configuration optimisation, neuromuscular coordination analysis, rehabilitation or FES-related applications, and modelling framework development. This focus aligns with traditional performance and injury questions. However, many studies lacked clearly articulated modelling goals or validation targets [23, 24, 26, 28, 31, 33, 48], echoing concerns that simulation research often begins with imprecise research questions [49]. Although internal variables such as joint loading and muscle forces were frequently estimated, few studies justified the specific joints, muscles or metrics chosen, and most did not model fatigue, metabolic cost or neural adaptation despite their known importance [4]. Neuromuscular coordination studies also tended to rely on simplified control assumptions rather than physiologically informed strategies, limiting their ability to probe muscle recruitment mechanisms.

Rehabilitation-focused studies demonstrated both promise and clear gaps. Simulations were applied to explore stimulation strategies [38], evaluate orthotic devices [39], and estimate joint loading in clinical cohorts [40]. Yet, only Farhoud et al. (2021) [38] implemented predictive optimal control, and none implemented personalised neuromuscular models or real-time applicability. This mirrors the gaps highlighted in [9, 50], which notes that current practices remain far from integration into rehabilitation planning, stimulation optimisation or patient-specific decision support. Even studies centred on SCI [38–40] or altered-gravity training [41] produced mainly descriptive insights without actionable recommendations. Overall, although methodological sophistication is increasing, the field has yet to address key translational needs such as clearer research questions, physiologically realistic control and subject-specific modelling.

Cycling protocols reinforced this pattern. Simulation-only studies [26, 31, 32, 37] typically used low cadences and low power outputs, likely reflecting intentional simplification for rehabilitation scenarios or limitations of existing forward and optimal control models. In contrast, experimental data were collected across a much broader range of intensities and setups, from instrumented road bikes to low-cost ergometers, showing that while data acquisition can accommodate diverse cycling conditions, simulation practices remain comparatively conservative.

Finally, participant characteristics revealed a clear demographic imbalance. Of the total individuals included, most were male, while relatively few were female (Fig 4(C)), and the majority were young, healthy, and recreational or trained cyclists. Clinical populations were only minimally represented (Fig 4(D)). Anthropometric ranges were similarly homogeneous. This pattern closely mirrors broader biomechanical bias reported in a recent meta-analysis [51], which identified pervasive male overrepresentation and limited justification for sex-imbalanced sampling. The alignment between our findings and field-wide evidence indicates that cycling biomechanics continues to rely on highly restricted populations, limiting generalisability and embedding the implicit assumption that young male morphology represents a default human model.

### Methodological frameworks

The musculoskeletal models used in cycling simulations varied widely in anatomical scope, dimensionality, muscle representations and degrees of freedom, ranging from single-leg 2D models to full-body 3D frameworks with up to 286 muscle–tendon units (Fig 4(E1) and (E2)). Yet key modelling details were often incomplete: nearly half of the studies did not report DOFs, and many omitted full muscle lists or failed to specify whether DOFs referred to the base human model (Table 4), the cycling-adapted configuration or the effective closed-chain system. These gaps reflect broader concerns raised in [10, 49], which note poor standardisation in reporting joint constraints, scaling procedures and muscle–tendon assumptions, as well as risks of unnecessary complexity without justification. Although model complexity varied greatly across studies, the rationale behind these choices was seldom articulated, limiting meaningful comparison of simulation behaviour.

Such variability also raises questions about task-specific suitability. Musculoskeletal models should be validated for the movement they simulate [52]. Yet, many cycling studies relied on gait-derived models, walking-optimised muscle parameters or knee formulations not validated for the large flexion ranges and closed-chain dynamics characteristic of pedalling (such as [53]). Most papers also used only basic geometric scaling, despite clear evidence that accurate muscle force and joint load estimation requires deeper personalisation, including subject-specific joint centres, muscle–tendon properties and neural control [9]. Limited personalisation, unclear constraints and sparse reporting therefore suggest that many simulations may not fully reflect cycling biomechanics, reducing confidence in internal load predictions.

Software choices reinforced these reproducibility challenges. Although OpenSim was the dominant platform (57.1%), followed by custom code (25.0%) and AnyBody (14.3%), only four studies (14.3%) shared models or code [39, 42, 43, 47] (Fig 4(F) and Table 5). This scarcity contrasts with long-standing recommendations to provide full documentation, parameter lists and accessible datasets [9], and with calls for open dissemination in ecosystems such as OpenSim, Moco and SCONE [49]. The predominance of open-source tools suggests these barriers are cultural rather than technical, echoing evaluations that biomechanical modelling must improve reporting completeness for reliable verification [10]. Overall, the field exhibits substantial reproducibility gaps despite the use of open-science platforms.

Across the included studies, data collection and validation practices were highly variable and often incomplete. Just over half collected new experimental data (53.6%), while the remaining studies relied on previously published datasets or conducted simulations without accompanying measurements. Although all data-based studies reported kinematics, far fewer included kinetics (69.6%), EMG (30.4%), or physiological signals (4.3%) (Fig 4(G) and Table 6), narrowing the scope for validation. Validation itself was inconsistent: three quarters compared simulations to experimental or literature data, yet most focused on kinematics or pedal forces, with only 66.7% evaluating muscle activation timing and none validating joint loads or resistance profiles. These patterns mirror broader challenges noted in [9], including that movement data reflect outcomes rather than control strategies and that EMG offers only qualitative validation, strongly influenced by normalisation and filtering that were often poorly documented. As emphasised in [52], many studies conflated verification with validation, providing little evidence that models were suitable for cycling-specific biomechanics. The absence of gold-standard internal load measurements, limited validation of stimulation-driven behaviour, and inconsistent reporting of optimisation settings further reflect the weak validation ecosystem described in [50]. Overall, while validation was frequently attempted, current practices remain insufficient to support strong confidence in simulated muscle forces, joint loads or neuromuscular behaviour.

Across the 28 studies, computational approaches ranged from static optimisation (32.1%) to forward dynamics and optimal control (each 17.9%), with inverse kinematics, computed muscle control and hybrid methods far less common (Fig 4(I1) and Table 7). This reliance on inverse methods reflects a broader tendency to analyse measured pedalling rather than predicting alternatives, consistent with the distinction in [49]: inverse simulations explain what occurred, whereas forward and optimal control explore why or how. Only a small subset of studies implemented fully predictive optimal control simulations, despite their potential for assessing equipment configurations, coordination strategies, or FES-driven pedalling. Limited adoption echoes challenges noted in [9], such as computational cost, solver sensitivity and dependence on accurate model parameters. Finally, none of the studies employed emerging approaches, such as real-time inverse dynamics, IMU, or video-based pipelines or ML-augmented methods, despite their growing relevance for out-of-lab analysis and digital twins. As highlighted in [10], the field is moving toward real-time control and personalised simulation, yet cycling research remains anchored to inverse pipelines that restrict predictive and translational potential.

Optimisation strategies showed similarly conservative tendencies. Among studies using explicit cost functions, 77.3% minimised muscle activity or effort, while few incorporated tracking, performance-related goals or physiologically grounded terms such as metabolic energy or joint loading (Fig 4(I2) and Table 7). This reliance on minimal-effort costs follows long-standing assumptions about cost-effective recruitment, but limits the exploration of alternative coordination strategies or fatigue-aware behaviours. For instance, FES-control research often relies on tracking-based formulations due to externally imposed movement goals [50], whereas the cycling studies used tracking terms far less often and mainly as secondary constraints. Yet, similarly to what was observed in FES [50], very few studies implemented multi-objective formulations balancing task accuracy with physiological realism, and almost none included fatigue despite its relevance for natural and electrically stimulated pedalling. Consequently, current optimisation practices prioritise movement replication over the discovery of more efficient biomechanical strategies.

Parameter variation was common: 27 of 28 studies modified at least one model, task or cost function parameter (Table 8), but these perturbations were mostly exploratory rather than structured sensitivity analyses. Most variations targeted geometric or equipment parameters (32.1%), whereas workload changes (21.4%) and musculoskeletal or morphological perturbations (21.4%) were less frequent. Only a small number of optimal control studies tested sensitivity to cost function weights or solver settings. In addition, the influence of different initial guesses on optimal control solutions was rarely examined, despite its importance for determining whether the solution obtained is a global optimum rather than a local one. These analyses also seldom quantified uncertainty or robustness, even though small changes in muscle–tendon parameters, joint definitions, or pedal constraints can markedly affect predicted forces and joint loads [9]. Known error sources in inverse simulations, including residual forces and parameter mis-specification, further emphasise the need for systematic sensitivity evaluation [49]. Overall, although parameter variation is widely used, the field lacks rigorous methods for uncertainty quantification, limiting confidence in the robustness and generalisability of cycling simulation outcomes.

### Limitations

Several limitations of this review should be acknowledged. First, the synthesis relied entirely on information reported in the included studies. Because many papers provided incomplete or inconsistent descriptions of modelling methods, data collection procedures, and validation approaches, some interpretation was required, and the extracted information may not fully reflect the authors’ original intentions. Second, the review included only peer-reviewed journal articles in English, potentially excluding relevant conference proceedings, theses, non-English publications, or unpublished modelling frameworks commonly shared within the biomechanics community. Third, the substantial heterogeneity of modelling approaches—spanning different software platforms, anatomical scopes, control methods, cost functions, and cycling protocols—prevented any form of quantitative synthesis or direct comparison of simulation outcomes. As a result, this review can characterise methodological trends and practices, but cannot evaluate the relative accuracy, performance, or clinical relevance of different modelling strategies.

### Future directions

The findings of this review highlight clear opportunities for the next generation of cycling simulation research. The field would benefit from greater methodological standardisation, deeper personalisation, and more transparent reporting, which would also create fertile ground for new work. Future studies could systematically test how model choices, DOFs, cost functions and scaling procedures influence predicted loads and coordination, helping establish cycling-specific best practices rather than relying on gait conventions. The strong demographic bias across existing datasets emphasises the need for more inclusive sampling and sex-specific modelling to examine how morphological and neuromuscular differences affect pedalling. Computationally, fully predictive optimal control simulations, real-time estimation pipelines, and wearable or machine-learning-enhanced approaches remain underused and could enable digital twin models for rehabilitation planning, equipment prescription, or adaptive FES control. Finally, formal uncertainty quantification and sensitivity analyses would help translate simulation outputs into clinically or performance-relevant decisions. By mapping current practices and limitations, this review lays the groundwork for more robust, generalisable and personalised cycling simulations that can advance both scientific insight and applied biomechanics.

## Conclusion

This review addressed two central questions regarding the current state of musculoskeletal cycling simulations, and the findings clarify both current practice and future needs. With respect to the first question, we showed that simulations are primarily used for equipment optimisation, neuromuscular coordination analysis, and rehabilitation applications. However, their scope remains limited by narrow participant profiles, conservative experimental protocols, and restricted demographic diversity, all of which constrain the generalisability of results. Regarding the second question, our synthesis revealed substantial variability across modelling frameworks, degrees of freedom, cost functions, and validation strategies. This variability highlights a lack of standardisation and reproducibility, while also identifying which methodological practices are well established, emerging, and underdeveloped. These conclusions are supported by a systematic methodology that adheres to transparent reporting guidelines, clearly defined inclusion criteria, and consistent data extraction across the 28 included studies. By integrating our findings with insights from key methodological literature, we situate current approaches within the broader landscape of musculoskeletal modelling and simulation. Together, these results provide a timely characterisation of the field, identify the methodological gaps that most limit interpretability and translation, and offer a foundation for more rigorous and personalised cycling simulations.

## Supporting information

**Supplementary material S1. Database search strategies** The following search queries were used to retrieve studies from each database.

### Scopus

~~~
TITLE-ABS-KEY({cycling} OR {pedalling} OR {pedaling} OR bicycle OR tricycle OR bike OR trike)
AND TITLE-ABS-KEY(biomechanic* OR neuromusculoskeletal OR neuromuscular OR musculoskeletal OR skeletal OR modeling)
AND TITLE-ABS-KEY(simulation* OR kinematics OR kinetics OR dynamics OR optimisation OR optimal OR tracking OR predictive
OR “machine learning” OR “artificial intelligence” OR “neural networks”) AND TITLE-ABS-KEY((joint W/15 (contact OR force OR position))
OR (muscle W/15 (effort OR force OR activity))
OR ((cycling OR pedaling OR crankset OR crank OR pedal) W/15
(cadence OR rate OR speed OR torque OR velocity OR power OR angle OR force)) OR (“muscle tendon unit”))
AND TITLE-ABS-KEY(participant OR human OR cyclist OR rider OR individual OR volunteer OR person
OR subject OR biker OR pilot OR children OR pediatric)
~~~

### IEEE Xplore

~~~
((((“Abstract”:cycling) OR (“Abstract”:pedalling) OR (“Abstract”:pedaling) OR (“Abstract”:bicycle) OR (“Abstract”:tricycle) OR (“Abstract”:bike) OR (“Abstract”:trike))
AND (((“Abstract”:biomechanic*) OR (“Abstract”:neuromusculoskeletal)
OR (“Abstract”:neuromuscular) OR (“Abstract”:musculoskeletal)
OR (“Abstract”:skeletal) OR (“Abstract”:modeling)))
AND (((“Abstract”:simulation*) OR (“Abstract”:kinematics) OR (“Abstract”: kinetics)
OR (“Abstract”:dynamics) OR (“Abstract”:optimisation) OR (“Abstract”:optimal) OR (“Abstract”:tracking) OR (“Abstract”:predictive)
OR (“Abstract”:“machine learning”) OR (“Abstract”:“artificial intelligence”) OR (“Abstract”:“neural networks”)))
AND (((“Abstract”:joint) NEAR/15 ((“Abstract”:contact) OR (“Abstract”:force) OR (“Abstract”:position)))
OR ((“Abstract”:joint) NEAR/15 ((“Abstract”:effort) OR (“Abstract”:force) OR (“Abstract”:activity)))
OR (((“Abstract”:cycling) OR (“Abstract”:pedaling) OR (“Abstract”:crankset)
OR (“Abstract”:crank) OR (“Abstract“:pedal)) NEAR/15
((“Abstract”:cadence) OR (“Abstract”:rate) OR (“Abstract”:speed)
OR (“Abstract”:torque) OR (“Abstract”:velocity) OR (“Abstract“:power)
OR (“Abstract“:angle) OR (“Abstract“:force)))
OR ((“Abstract“:”muscle tendon unit“)))
AND ((“Abstract”:participant) OR (“Abstract”:human) OR (“Abstract”:cyclist)
OR (“Abstract“:rider) OR (“Abstract”:individual) OR (“Abstract”:volunteer)
OR (“Abstract“:person) OR (“Abstract”:subject) OR (“Abstract”:biker)
OR (“Abstract“:pilot) OR (“Abstract”:children) OR (“Abstract”:pediatric)))
~~~

### PubMed

~~~
((”cycling“[Title/Abstract]) OR (pedalling[Title/Abstract]) OR (pedaling[Title/ Abstract])
OR (bicycle[Title/Abstract]) OR (tricycle[Title/Abstract])
OR (bike[Title/Abstract]) OR (trike[Title/Abstract]))
AND ((biomechanic*[Title/Abstract]) OR (neuromusculoskeletal[Title/Abstract])
OR (neuromuscular[Title/Abstract]) OR (musculoskeletal[Title/Abstract])
OR (skeletal[Title/Abstract])
OR (modeling[Title/Abstract]) OR (modelling[Title/Abstract]))
AND ((simulation*[Title/Abstract]) OR (kinematics[Title/Abstract])
OR (kinetics[Title/Abstract]) OR (dynamics[Title/Abstract])
OR (optimisation[Title/Abstract]) OR (optimal[Title/Abstract])
OR (tracking[Title/Abstract]) OR (predictive[Title/Abstract])
OR (“machine learning”[Title/Abstract])
OR (“artificial intelligence”[Title/Abstract])
OR (“neural networks”[Title/Abstract]))
AND ((participant[Title/Abstract]) OR (human[Title/Abstract])
OR (cyclist[Title/Abstract]) OR (rider[Title/Abstract])
OR (individual[Title/Abstract]) OR (volunteer[Title/Abstract])
OR (person[Title/Abstract]) OR (subject[Title/Abstract])
OR (biker[Title/Abstract]) OR (pilot[Title/Abstract])
OR (children[Title/Abstract]) OR (pediatric[Title/Abstract]))
AND (((joint[Title/Abstract]) AND ((contact[Title/Abstract])
OR (force[Title/Abstract]) OR (position[Title/Abstract])))
OR ((joint[Title/Abstract]) AND ((effort[Title/Abstract])
OR (force[Title/Abstract]) OR (activity[Title/Abstract])))
OR (((cycling[Title/Abstract]) OR (pedaling[Title/Abstract])
OR (crankset[Title/Abstract]) OR (crank[Title/Abstract])
OR (pedal[Title/Abstract])) AND ((cadence[Title/Abstract])
OR (rate[Title/Abstract]) OR (speed[Title/Abstract])
OR (torque[Title/Abstract]) OR (velocity[Title/Abstract])
OR (power[Title/Abstract]) OR (angle[Title/Abstract])))
OR (“muscle tendon unit”[Title/Abstract]))
~~~

### Web of Science

~~~
#1: AB=(cycling OR pedalling OR pedaling OR bicycle* OR tricycle* OR bike* OR trike*)
#2: AB=(biomechanic* OR neuromusculoskeletal OR neuromuscular OR musculoskeletal OR skeletal OR modeling OR modelling)
#3: AB=(simulation* OR kinematic* OR kinetic* OR dynamic*
OR optimisation OR optimal OR tracking OR predictive
OR “machine learning” OR “artificial intelligence” OR “neural networks“)
#4: AB=((joint* NEAR/15 contact) OR (joint* NEAR/15 force*)
OR (joint* NEAR/15 position)
OR (muscle* NEAR/15 (effort OR force* OR activity))
OR ((cycling OR pedaling OR crankset OR crank OR pedal)
NEAR/15 (cadence* OR rate OR speed* OR torque*
OR velocity OR power OR angle* OR force*))
OR (“muscle tendon unit”))
#5: AB=(participant* OR human* OR cyclist* OR rider*
OR individual* OR volunteer* OR person*
OR subject* OR biker* OR pilot* OR children OR pediatric OR child)
(#1 AND #2 AND #3 AND #4 AND #5)
~~~

## Notes

### Competing Interest Statement

The authors have declared no competing interest.

